# Primary infection by *E. multilocularis* induces distinct patterns of crosstalk between hepatic regulatory T and natural killer T cells in mice

**DOI:** 10.1101/2021.12.08.471874

**Authors:** Tural Yarahmadov, Junhua Wang, Daniel Sanchez-Taltavull, Cristian A. Alvarez Rojas, Tess Brodie, Isabel Büchi, Adrian Keogh, Bruno Gottstein, Deborah Stroka, Guido Beldi

**Affiliations:** Department of Visceral Surgery and Medicine, Inselspital, Bern University Hospital, University of Bern, Switzerland; Institute for Infectious Diseases, University of Bern, Bern, Switzerland; Institute of Parasitology, University of Zurich, Zurich, Switzerland

**Keywords:** *Echinococcus mutilocularis*, single-cell RNA sequencing, Tregs, NKT cells

## Abstract

The larval stage of the helminthic cestode *Echinococcus multilocularis* can inflict tumor-like hepatic lesions that cause the parasitic disease alveolar echinococcosis in humans, with high mortality in untreated patients. Recently, opportunistic properties of the disease have been proposed based on the increased incidence in immunocompromised patients and mouse models, indicating that an appropriate adaptive immune response is required for the control of the disease. However, little is known about how the local hepatic immune responses modulate the infection with *E. multilocularis*. In a mouse model of oral infection that mimics the normal infection route in human patients, the adaptive immune response in the liver was assessed using single-cell RNA sequencing of isolated hepatic CD3^+^ T cells at different infection stages. We observed an early significant increase in regulatory T and natural killer T cells in parallel with an active downregulation of CD4^+^ and CD8^+^ T cells. Early interactions between regulatory T cells and natural killer T cells indicate a promotion of the formation of hepatic lesions and later contribute to suppression of the resolution of parasite-induced pathology. The obtained data provides a fresh insight on the adaptive immune responses and local regulatory pathways at different infection stages of *E. multilocularis* in mice.

**Author summary:** Alveolar echinococcosis is an endemic parasitic infection leading to slowly growing but potentially lethal liver lesions if untreated. Transmission by increasing populations of urban foxes and the raise of immunosuppressed patients in mainly industrializsed endemic zones are the main causes responsible for increased incidence of alveolar echinococcosis. Observations in humans and mice indicate that reactions of the adaptive immune system are required to control the disease and to protect from a chronic infection. Therefore, we analysed the responses of T cells in the liver at single-cell resolution in a murine model mimicking the typical route of infection in humans. This so called single-cell RNA sequencing revealed specific temporal changes of T cell subsets such as natural killer T cells and regulatory T cells, indicating that these two cell types are recruited in the early phase to try to protect from parasite proliferation and are subsequently inhibited in the late phase of infection thus indicating immune escape mechanisms of the parasite. This study shows temporal changes of the immune cell profile in the liver over the course of a natural infection with *E. multilocularis* at the single cell level and reveals putative targets for novel therapeutic approaches for human AE and possibly other (parasitic/helminthic) diseases.

## Introduction

Alveolar echinococcosis (AE) is a rare but lethal zoonotic helminthic disease if remained untreated, causing direct damage to the liver by the parasitic infiltration and mass proliferation and indirect damage by modulating the periparasitic host response. The continuous proliferation of the larval stage (metacestode) of *E.multilocularis* leads to an intense local granulomatous immune response surrounding the parasitic tissue thereby leading to tumor-like focal lesions [1,2].

Clinical observations revealed an elevated incidence of AE in immune compromised patients indicating that the incidence and the course of the disease may depend on the immune response of the host [3]. The increase of case numbers in the last decades can therefore be explained in part due to the increased number of patients receiving immunosuppressive therapy, in addition to the increased fox population density and associated environmental contamination with parasite eggs [4–6]. Therefore, understanding the immune response to AE is required to apply potential immunomodulatory treatments.

Previous studies on the interaction between the parasite and the host immune system were performed in immune compromised mice and in humans [7–10]. Initial parasite survival and proliferation correlated with a Th2-reorientation of the CD4^+^ T lymphocytes, leading to immunotolerance or to an anergic response to infection at later stages [1,8,11,12]. Resistance of the parasite was associated with Th1-cytokine profiles at later stages including IFN-α [13] and IL-12 [14] as initiating cytokines, and IFN-γ [15] and TNF-α [16,17] as effector cytokines. Furthermore, AE-patients with aborted lesions presented lower secretion levels of IL-10 and Th2 cytokines by PBMCs than patients with a progressing disease [13]. Thus, studies have shown that AE is more severe, with a faster course, when there is some degree of impairment of functional activity of T cells and persistence of *E. multilocularis* is associated with a chronic granulomatous inflammation, leading to the disruption of the normal function of T cells which is referred to as ‘functional exhaustion’ [18].

These studies are limited as the most conventional infection model applied resembles secondary AE, i.e. infection induced upon intraperitoneal inoculation of *E. multilocularis* metacestode [19,20]. The problem associated to this infection model is that it lacks the first phase of passage of the parasite through the intestinal barrier and the second phase of intrahepatic parasite development both being potentially crucial for the formation of liver lesions [21,22]. Therefore, a model of primary infection using orally ingested tapeworm eggs seems to be preferential as it is representative of the natural route of infection. The few studies using a primary infection model, have described a systemic humoral response when the metacestode invades the liver [23,24] and a tolerogenic state at the late stage of AE [20].

Thus far, no study has reported on the changes of *in situ* host T-cell immune responses from early to chronic stage of primary murine AE. For the first time, we used single-cell RNA sequencing in a murine model of primary AE having the capabilities to identify transcriptional changes at the individual cellular level to identify disease specific alterations. Given the relevance of adaptive immune responses in AE, the aim of this study was thus to give a comprehensive analysis of the different T cells involved in immune cell homing at the various successive stages of disease, i.e., early, intermediate, and chronic stages. Therefore, CD3^+^ T cells were isolated from the liver at 10, 21, and 48 days post primary infection, and the T cell immune response was assessed by using single-cell RNA sequencing. Analysis revealed novel temporal changes of regulatory T cells (Treg) and natural killer T cells (NKT)and associated pathways indicating yet unknown functions involved in disease pathology.

## Results

### Expansion of natural killer T cells and regulatory T cells as a first response to infection

We analyzed a dataset spanning over 4 time points pre and post oral infection. The dataset consists of hepatic T cells from non-infected control mice (control: D0, 2 samples), and the cells from livers of mice at 10 days (early infection: D10, 2 samples), 21 days (intermediate infection: D21, 3 samples), and 48 days (late infection: D48, 3 samples) post oral infection (Figure 1A). Together, the studied dataset consists of 14848 cells from 10 different samples. Unsupervised clustering using Seurat was performed and manually adjusted based on higher clustering resolution level to better separate immune cell subpopulations (see Methods section “Dimensionality reduction and filtering”) (Figure 1B, C). Cell types were next annotated using SingleR [25] (Figure 1D, E, Figure S1), resulting in 15 clusters overlaid with the predicted cell annotations showing spatial separation of the different cell types.

**Figure 1.**
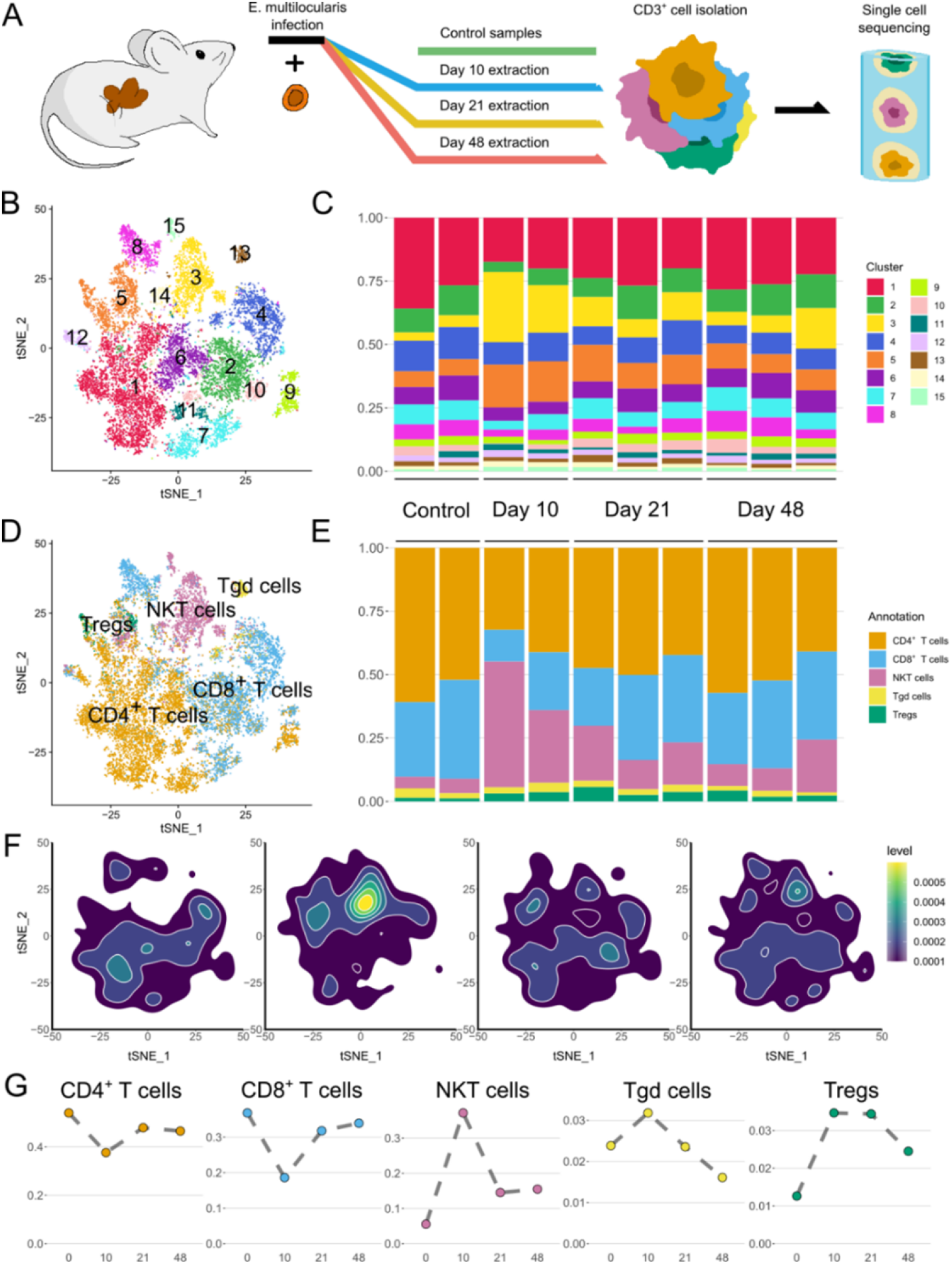
Clustering of the dataset spatially and temporarily separates cell types. **A** – Schematic representation of the experiment. CD3^+^ cells from the liver of control mice and mice at 10, 21, and 48 days post peroral *E. multilocularis* infection were extracted for single-cell RNA sequencing. **B** – t-distributed stochastic neighbor embedding (tSNE) of the final dataset based on the single-cell RNA-seq data coloured by clusters. **C** – Normalized distribution of cell subsets from each cluster in samples are displayed in chronological order from left to right. **D** – tSNE of the annotation using SingleR to represent major T cell types in the dataset. **E** – Normalized cell numbers (cell subset distributions) from each cell type in chronological order. **F** – Density maps of cells at different stages. Highest quantity of NKT cells can be observed at D10 compared to the rest. **G** – Distribution of relative to the total dataset number of cells (Y axis) over the course of the experiment (X axis) for all cell types.

According to SingleR annotation, the final dataset consists of 7057 CD4^+^ T cells (48%), 4805 CD8^+^ T cells (32%), 2297 natural killer T cells (NKT, 15 %), 372 regulatory T cells (Tregs, 3%), and 317 T gamma delta cells (Tgd, 2%) (Table S1). To identify cell types that were differentially abundant over the course of the experiment, we quantified the number of cells belonging to different types at each time point. Then, we tested them by fitting the values to quasi-likelihood negative binomial generalized log-linear model, producing p-values adjusted by Benjamini-Hochberg method or false discovery rate (FDR, for details see “Differential abundance testing” section of the Methods). At D10 NKT cells expanded sharply (FDR 0.0004) and to a lesser extent also the abundance of Tregs increased (FDR 0.043) (Figure 1D, F, G). No statistical significance was observed for other cell types or time points.

### Unbiased clustering identifies early downregulation of CD4^+^ and CD8^+^ cells

To better understand the assembly and dynamics of the transcriptome of the cellular subsets we next performed unbiased differential abundance testing of cellular subpopulations (Figure 2A). Cell numbers at each time point were compared to control samples only and to the average of the other time points, and consecutive time pointwise (Figure 2B). Significant changes were found for clusters 2, 3, 5, 6, 10, and 15 at D10. The clusters 3 and 15 elevated at D10 consist predominantly of NKT cells while cluster 5 contains the majority of Tregs. Clusters 2, 6, and 10 decreased at D10 consist mostly of CD4^+^ and CD8^+^ cells at D10 post oral infection with *E. multilocularis* eggs as shown in Figure 1G (Figure 2C, Table S1). The majority of all NKT cells in the dataset are represented in cluster 3 (1491/2297, 65%) and to a lesser extent in cluster 15 (95%) and conversely cluster 3 consists almost entirely of NKT cells (1491/1573, 95%) (Figure 2C). This observation is further supported by the expression e.g. of Zfp683/Hobit in this cluster, which is a gene important for proper NKT cell development and morphology [26] (Figure 2D). Cluster 15 exhibits markers characteristic of both NKT and NK cells, indicating some potential remaining subsets of NK cells in the dataset after filtering. The majority of Tregs in the dataset are represented in Cluster 5. This cluster is heterogenous and is split between 667 CD4^+^ T cells (48%), 323 Tregs (23%), 267 NKT cells (19%), and 125 CD8^+^ T cells (9%) (Figure 2B, C; Table S1). CD8^+^ T and CD4^+^ T cells are represented in Cluster 2, 6 and 10. Cluster 2 is split between 1243 CD8^+^ T cells (75%) and 398 CD4^+^ T cells (24%) (Figure 2B). It displays high expression of CD8B1, a cell surface glycoprotein characteristic of cytotoxic CD8^+^ T lymphocytes [27]. Cluster 6 contains 871 CD4^+^ T cells (69%) and 345 CD8^+^ T cells (27%) (Figure 2B) and is characterized by the high expression of long non-coding RNAs Gm42418 and Gm26917 (Figure 2D). Cluster 10 is evenly divided between 221 CD8^+^ T cells (53%) and 191 CD4^+^ T cells (46%) (Figure 2B). This cluster is also remarkable for its strong and almost exclusive caspase 11 (Scaf11) expression (Figure S3). Interestingly, the two clusters, 2 and 10, underrepresented at D10 are among the few remarkable for strong expression of genes encoding ribosomal proteins indicating an active downregulation of the associated CD4^+^ and CD8^+^ T cells (Figure S4) thereby representing another mechanisms that decrease of CD4^+^ and CD8^+^ T cells in addition to the induction of pyroptosis via capsase-11. These unbiased data support the findings of Figure 1 showing mechanisms of the elevation of NKT and Treg and an active downregulation of CD4^+^ and CD8^+^ cells at D10.

**Figure 2.**
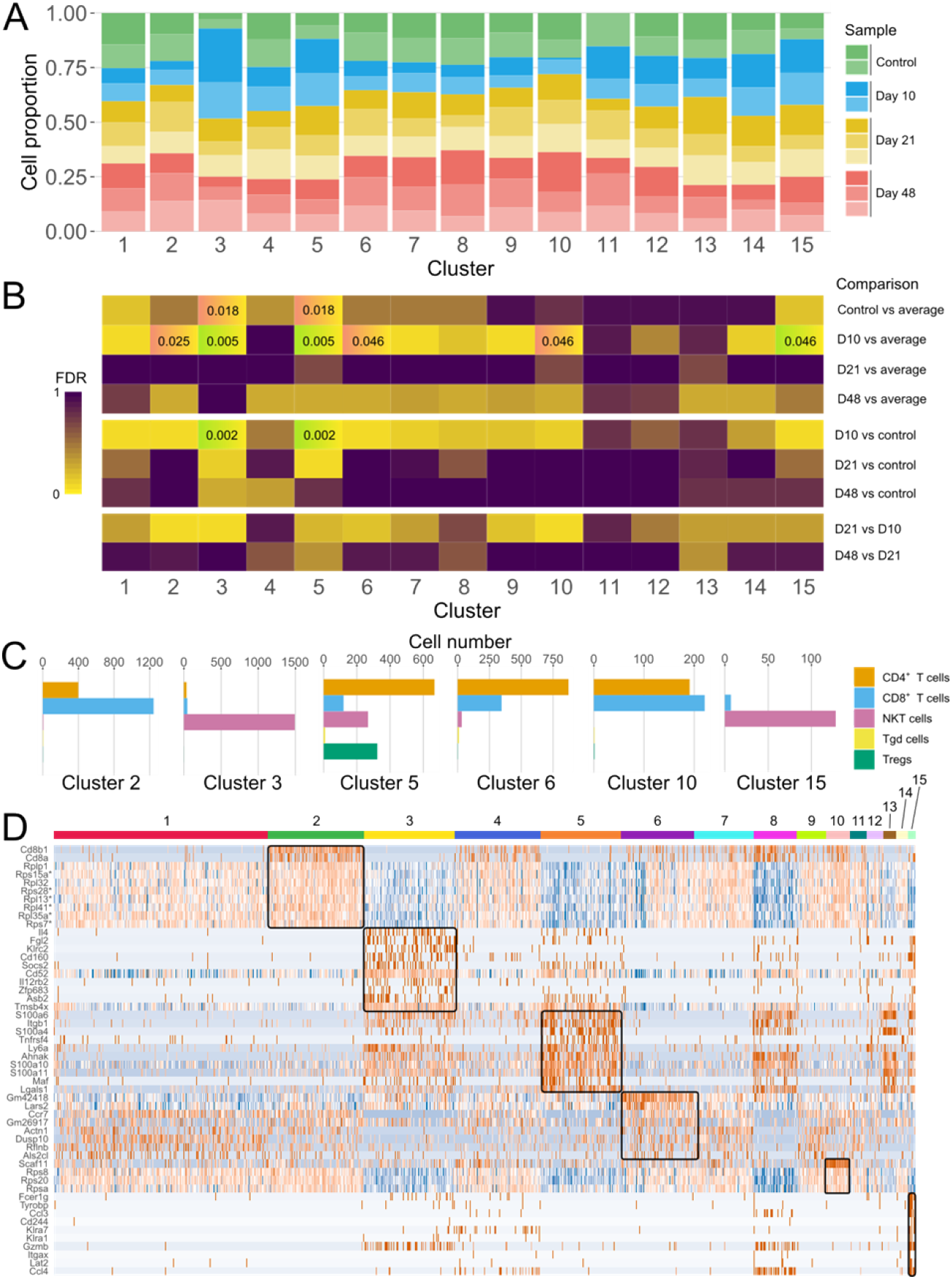
Cluster distribution of samples shows temporal dynamics of cluster sizes over time of the infection. **A** – Distribution of sample cell numbers in each cluster over time. Normalized cell numbers from each sample are displayed per cluster. **B** – Results of differential abundance testing of number of cells in each cluster compared to the average number across other time points. Cells with values of FDR < 0.05 are labelled and tinted in green in case of higher number and red of lower number, respectively. **C** – The majority of NKT cells and Tregs is contained in overrepresented D10 clusters. Underrepresented clusters contain only CD4^+^ and CD8^+^ cells. **D** – Heatmap of most differentially expressed marker genes from clusters of interest. Ten most significant markers by p-values and log2fold change per cluster are depicted. Each line represents expression value in one cell. Expression values are Z-score transformed by row, with low values being blue, intermediate – white, and high – red. Genes and cells from differentially abundant clusters are highlighted via rectangles. *Highlighted markers are shared between clusters 2 and 10.

### Interactions between NKT and Treg mediate early recruitment and regulation of the responses at late phases

After having identified genes associated with expansion and activation of NKT and Treg cells at D10, we aimed to analyze the cross-talk between different immune cell types and subtypes (Figure S5). We performed a receptor-ligand complex analysis via CellPhoneDB, using mouse orthologs from ENSEMBL to access the human database (Figure 3) [28,29]. A consistently high number of interactions between Tregs, NKT and NK cells was observed, while CD4^+^ T and CD8^+^ T cells showed low amounts of predicted interactions between themselves and each other (Figure 3A-E, Figure S6). Most of the interaction pairs also display a reduction in number at D10 with subsequent return to the baseline number of connections to other cell types over the course of the experiment (Figure 3E). This pattern is also observed in pairs formed by NKT cells and Tregs, which expand at D10 indicating a transient suppression of cellular interactions at this time point (Figure 1G). While the number of predicted interactions between Tregs and NKT cells remains comparatively high throughout the experiment (Figure 3), the composition of interacting pairs changes across time points (Figure 4). Relevant interactions between NKT and Treg cells are shown in Figure 4. At early phases include regulation of cell adhesion molecules such as integrins and selectins and TNF-dependent signalling dictates the interactions between NKT and Treg. A strong upregulation of alb2 complex (LFA-1) occurs on NKT cells at D10 when also these cells are highly enriched in the course of infection. The upregulation of ICAM1 and alb2 complex at D10 occurs in parallel to the regulation of the selectin interaction SELPLG-SELL. At the late time point D48, expression of alb2 complex in NKT ceases conversely to Treg in which alb2 complex is highly upregulated. Immune suppressing interactions that include the checkpoint inhibitors (PD-1 and PD-L1), purinergic (ENTPD1, ADORA2A) and TGF-beta signalling are down regulated at D10 and D21 returning to baseline values at D48. In order to evaluate the connectivity between proteins forming the NKT cell and Tregs interaction pairs in control and D10 time points their protein-protein interaction (PPI) networks were built via STRING database interface (Figure S7). All resulting networks were found to be strongly interconnected with the number of edges being significantly higher than expected (PPI enrichment p-value < 1.0e-16). Networks made from proteins forming pairs in control samples have higher average degree than those made from D10 proteins in both NKT cells (7.65 to 3.89) and Tregs (7.85 to 4.95).

**Figure 3.**
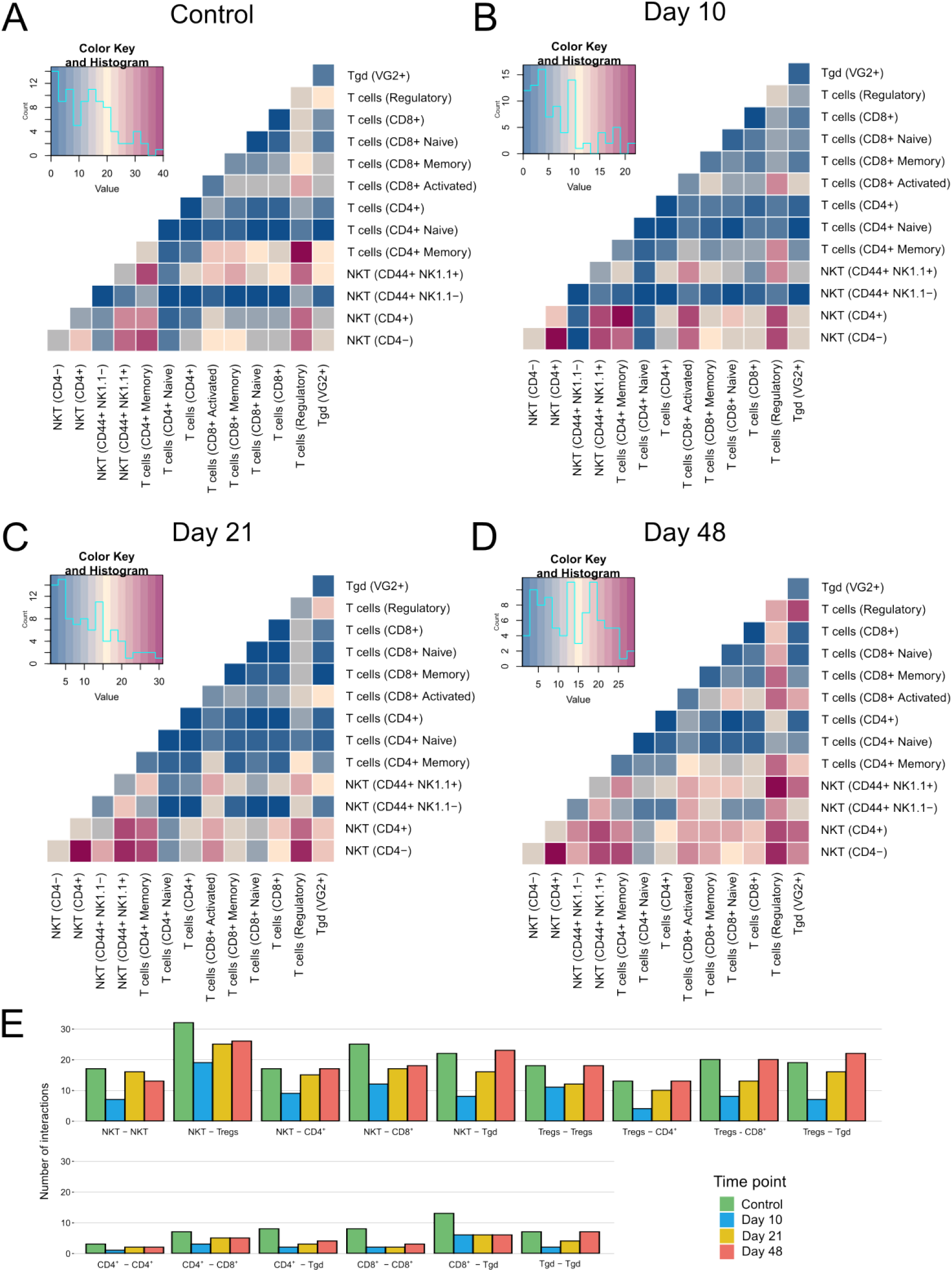
Maps of intercell interactions show a consistently high number of NKT cell and regulatory T cell interactions over the course of the experiment. Heatmaps summarizing statistically significant ligand-receptor interactions per cell type pair can be seen for control (**A**), D10 (**B**), D21 (**C**), and D48 samples (**D**). **E –** Total amount of significant interactions between pairs of cell types indicating a decrease at D10.

**Figure 4.**
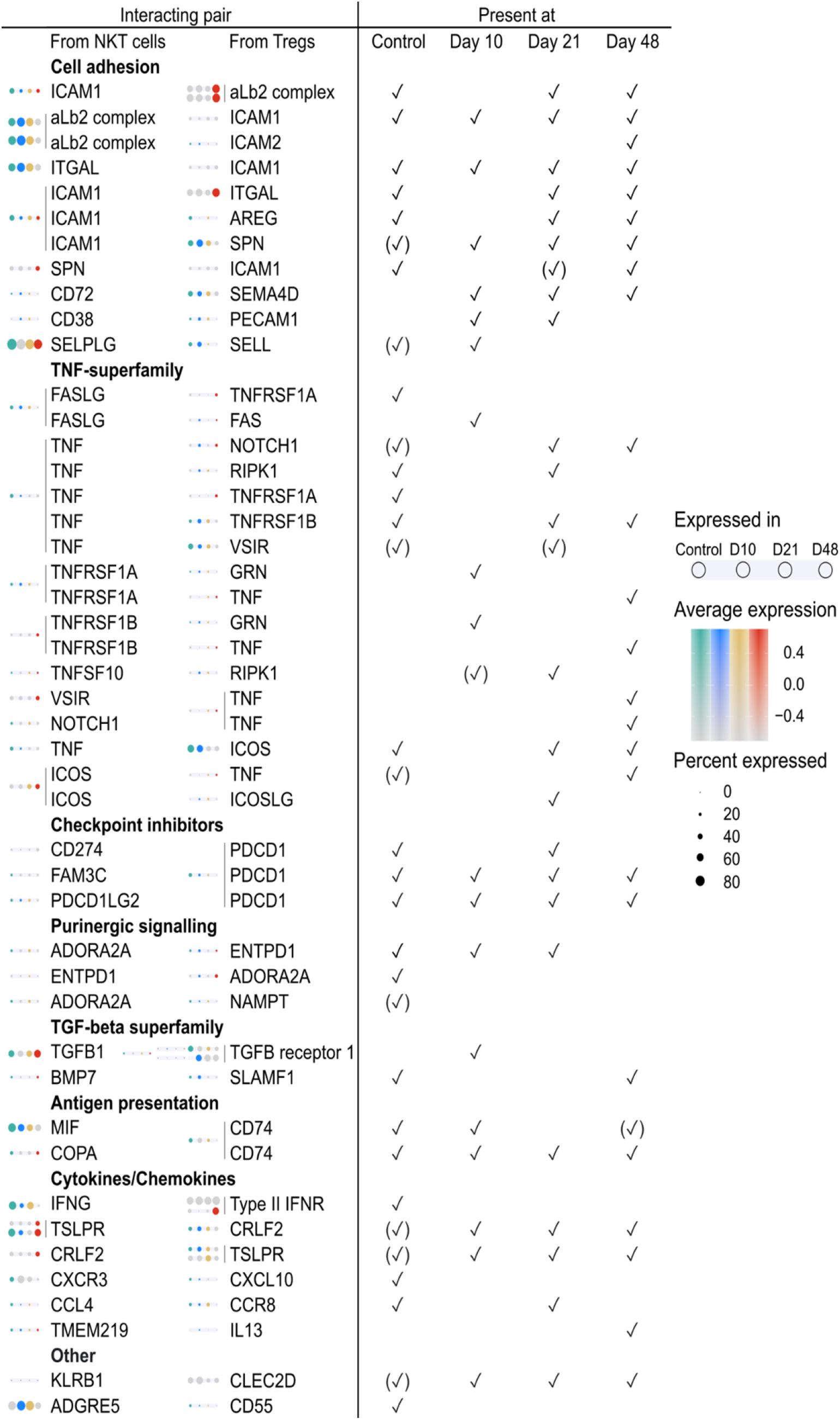
Genes associated with interactions between NKT cells and Tregs. Analysis a receptor-ligand complex analysis was performed via CellPhoneDB, using mouse orthologs from ENSEMBL to access the human database.

## Discussion

Based on studies in mice and humans, adaptive T cell-mediated responses dictate the outcome after infection respective to the development of AE [10]. In the present study, we investigated the *in situ* host T cell response involved in immune cell homing at the various successive stages of disease (early, middle and chronic stage) in a model of primary murine AE. By performing single-cell RNA sequencing of isolated CD3^+^ T cells, we showed a significant increase of NKT cells and to a lesser extent Treg cells at D10 post infection with a subsequent decrease at later time points (D21-48) to the levels of non-infected controls (Figure 1). This indicates a peak of the acute immune response to *E. multilocularis* infection around 10 days post oral infection, with a subsequent stabilization once the lesions were grossly visible in the liver (14 days post oral infection in mice, not shown).

Unbiased analysis of the data revealed 6 subclusters with significant differences in abundance over time (Figure 2). Consistent with alterations in cellular subsets (Figure 1), the differences were predominant at D10 with 3 clusters being over- and 3 under-represented at that stage. The over-represented clusters 3 and 15 underlines the relevance of NKT cell activity (Figure 3A, Figure 3B), which supports the role of NKT found in other in macroparasitic infections [30,31]. A gene important for proper NKT cell development and morphology, Zfp683/Hobit [26], is a strong marker for cluster 3, thus confirming the results of the automatic annotation of the individual cell subsets (Figure 2C). Our data are in contrast with previous findings which suggested NKT cells played a rather minor role against *E. multilocularis* [18]. These previous data are, however, derived from a model of secondary infection in which intestinal update and hepatic seeding of AE cannot be assessed. Our current study indicates that NKT cells possibly play an important role at early infection stage of primary AE, in which the natural course of the disease is modelled.

Treg cells also seem to be of relevance beginning at D10 and later time points (Figure 3C, 3D). Their cluster is highlighted by a comparatively high level of S100 protein family expression, namely S100a4, S100a6, S100a10, and S100a11 (Figure 2C, Table S2) promoting motility in T cells, cell proliferation, cytoskeleton formation and tumorigenesis [32–36]. Under homeostatic conditions, genes of the TNF superfamily such as the receptor TNFRSF4, which play a role in the activation of CD4^+^ and CD8^+^ T cells, as well as Tregs are strongly upregulated [37,38]. These data support the role of Treg which were identified in studies of AE patients [39] and in secondary experimental models [20,40,41]. Together these data indicate that metacestode persistence is the consequence of immune tolerance or anergy, respectively, mainly mediated by specialized Treg cells and related cytokines such as IL-10 and TGF-β [20].

Tregs and NKT cells are two populations of T lymphocytes that can independently regulate adaptive and innate immune responses. Recent reports provide evidence for cross-talk between Tregs and NKT cells including the ability of NKT cells to regulate the suppressive activity of Tregs [42] in anti-tumor immune responses [43], and in the prevention of autoimmune myasthenia [44]. With regards to microbe and parasite infections, especially at early lifetime, parasitic worms are able to survive in their mammalian host for many years due to their ability to manipulate the immune response [45]. Upregulation of regulatory T cell subsets, such as Treg, and induction of inhibitory cytokines and/or chemokines are the common findings [45]. The involvement of NKT cells in down-regulation of the immune responses mostly remains elusive. Given the fact that many microbes and parasites are enriched in lipid antigens and NKT cells are the unique T cell subset that can recognize lipid antigens, it is reasonable to speculate that NKT cells play key roles [46].

Analysis of interactions between NKT and Treg cells revealed upregulation of integrin signalling that seems to be associated with the development of the early disease. Conversely, genes associated with immune suppression such as checkpoint blockade (PD-1/PD-L1) or adenosine signalling (ENTPD1/ADORA2A) are upregulated early and downregulated at late phases, respectively. Downregulation of these genes at the late time points indicates a response of the local immune system to control the parasite. Potentially, such an autonomous checkpoint inhibition might be the cause of the intense immune reaction with dense fibrous walls around the parasite as observed in mice and humans.

The present data also reveal the decrease of different cellular subsets at D10. For example, it appears there is CD8^+^ T cell dependent cytotoxicity at D10. This is supported by the decrease in cell number in its associated cluster and that it exhibits high ribosomal gene activity indicating translational activity of the cells (Figure 2C, Figure S4) [27]. There is a reduction of caspase 11 (Scaf11) expressing CD4^+^ T and CD8^+^ T cells at D10 (Cluster 10, Figure 2C, Figure S3). The decreased number of cells with activated caspase 11 in our model may be the consequence of associated pyroptosis and cell death [47,48] that may be triggered by the parasite by yet unknown mechanisms. The under-represented CD4^+^/CD8^+^ cell cluster 6 can be characterized by high expression of long non-coding RNAs Gm42418 and Gm26917, which were found to be enriched in NLRP3 inflammasomes which also trigger inflammatory cell death and release of cytokines [49,50]. Causes and consequences of such downregulation of these subsets of CD4^+^ and CD8^+^ T cells at an early infection stage need to be further addressed in future studies.

Together, these comprehensive findings on the immunological changes at different stages in primary *E. multilocularis* infection now open the door for more applied approaches. For instance, the experimental infection of caspase11^-/-^’ mice or further addressing the role of checkpoint blockade or purinergic signalling could provide tools of converting the immunological anergy during chronic disease into a more pro-inflammatory, with potentially fatal consequences for the metacestode. Furthermore, our findings may provide a rationale for studying immune cells as a target for an immunomodulatory treatment option in patients with progressive AE.

## MATERIALS AND METHODS

### Animal studies

#### Ethics Statement

The animal studies were performed in accordance with the recommendations of the Swiss Guidelines for the Care and Use of Laboratory Animals. The protocol was approved by the governmental Commission for Animal Experimentation of the Canton of Bern (approval no. BE112/17).

#### Mice

36 Female 8-week-old wild type C57BL/6-mice were purchased from Charles River GmbH (Sulzfeld, Germany), and divided randomly into groups as follows: 1) *E. multilocularis* infected (named as “AE”); 2) non-infected control (named as “Control”) at three time points (D10, D21, D48 post oral infection, 6 mice per group/time point).

#### Parasite and experimental infection

*E. multilocularis* eggs were isolated from a naturally *E. multilocularis-infected* foxes, which was euthanized at the Institute of Parasitology, Vetsuisse Faculty in Zurich. Infection with *E. multilocularis* was detected upon routine necropsy investigation by pathologists. To prepare the parasite eggs for subsequent infection of mice, the fox intestine was removed under appropriate safety precautions and cut into 4 pieces. After longitudinal opening of the intestinal segments, the worm-containing mucus was scraped out and put into petri dishes containing sterile water. Subsequently, the mucosal suspension was serially filtered through a 500μm and then 250μm metal sieve, by concurrently disrupting the worms with an inversed 2 ml syringe top. This suspension was further filtered through a 105 μm nylon sieve. The eggs were then washed by repeated sedimentation (1xg, 30 min., room temperature) in sterile water containing 1% Penicillin/Streptomycin and stored in the same solution at 4 °C. For primary infection of mice, animals were received approximately 400 eggs suspended in 100 μL sterile water by peroral gavage. Control mice (mock-infection) received 100 μL water only. All animal infections were performed in a biosafety level 2plus unit (permit no. VTHa-R9).

#### Sampling

At control, D10, D21 and D48 post infection, mice were sacrificed by CO_2_-euthanasia. At necropsy, the number and size (diameter) of the individual liver lesions (each caused by one developing oncosphere, originating from one parasite egg) were recorded. Liver cells were isolated from 3 AE mice/time point with successful liver lesions by using percoll density gradient centrifugation (see below).

### Hepatic lymphocyte isolation

Mouse livers were cut into small pieces with a scalpel and crushed against a 50mm stainless steel mesh plate. The mesh plate was rinsed with cold MACS buffer (PBS with 0.6% FBS and 0.5M EDTA) to release the cells into the collection plate. Cells were washed with MACS buffer and centrifuged at 50G for 3 minutes at 4°C to get rid of most of the hepatocytes and the supernatant was collected (non-parenchymal cells). The non-parenchymal cells were then centrifuged at 300G for 7 minutes at 4°C. The pellet was resuspended in 10mL of 40% Percoll and overlaid with 70% Percoll. The cells were centrifuged at 800G for 20 minutes at 4°C with no brake and the interphase with cells were collected and resuspended up to 15mL in MACS buffer. The cells were centrifuged at 300G for 7 minutes at 4°C and the cell pellet was resuspended then counted with the BioRad cell counter (Bio-Rad, TC20).

### CD3 negative selection

0.25-0.5 million non-parenchymal cells were used with the EasySepTM Mouse T cell Isolation Kit (Stemcell Technologies, Cat.# 19851) for negative selection of T cells, following kit protocol. Briefly, cells are stained with a cocktail of antibodies bound to magnetic particles and specific for non-T cells. These cells are retained against the wall of the FACS tube when the EasySep magnet is applied and the remaining cells (T cells) are poured out of the tube and counted for sequencing.

### Sequencing

Between 5,000 and 20,000 cells per sample were provided to the Next Generation Sequencing Platform at the University of Bern and standard methods were used to perform the sequencing. Read processing and data processing steps were performed with the help of UBELIX (http://www.id.unibe.ch/hpc), the HPC cluster at the University of Bern.

### Read processing

Feature barcode counting was performed using 10x Genomics cellranger v.3.0.2 *count* function with software-provided mouse reference genome mm10-3.0.0 and –*expect-cells* parameter value 5000.

### Data processing

Obtained data was analysed in R v.3.6.1 [51]. Data was imported into R and analysed using Seurat R package v. 3.1.4 [52]. Filtered count matrices produced by cellranger *count* function per sample were imported into R as Seurat objects using *Read10X* function. Cells in samples were individually filtered based on gene (feature) content and proportion of mitochondrial mRNA by Seurat *subset* command. For sample C1D10 the threshold was to have at least 100 genes, for sample AE2D10 – 150 genes, for sample AE3D10 – 200 genes, for samples AE1D48, AE2D48, C1D21, C1D48 – 250 genes and for samples AE5D21, AE6D21, AE7D21, AE3D48 – 300 genes. All samples were filtered to have cells with at most 15% of mitochondrial RNA. All samples were combined using Seurat function *merge*. Samples AE3D10 and C1D10 were excluded based on initial clustering analysis due to abnormal clustering and high mitochondrial content. Remaining samples were merged into single object by applying Seurat *SCTransform* based on 3000 anchoring features with parameter *vars.to.regress = “percent.mt”*.

### Cell annotation and filtering

Data was analysed with seed 322. Cells were annotated using immgen database (02-2020) [53] with the help of SingleR R package v.3.10 [25], “main” and “fine” annotations. Only cells with related to CD3^+^ T cells annotations - having “T cells”, “Tgd”, and “NKT” - were kept for further analysis (15228 kept out of 16327, 6% discarded). CD4+ and CD8+ T cells were distinguished based on “fine” level annotations.

### Dimensionality reduction and clustering

Principal component analysis (PCA) was performed on the full post-filtered dataset using Seurat *RunPCA* function. A UMAP dimension reduction was performed using Seurat *FindNeighbors* and *RunUMAP* functions with parameter *dims = 1:24*, which was chosen based on *ElbowPlot* function graph. Clusters were defined via *FindClusters* function at resolution 0.3.

### Data filtering and reclustering

Positive cluster markers were identified by *FindAllMarkers* function with parameters *only.pos = T, min.pct = 0.25, logfc.threshold = 0.25*. Cell cycle stage of cells was assessed with function *CellCycleScoring*, with *s.features* and *g2m.features* provided by Seurat in the R object *cc.genes* after being transformed into mice genes with the R function *gorth* of the R package gprofiler [54]. Based on marker and cell cycle analysis, clusters 14 and 16 of likely actively proliferating cells were discarded. Remaining cells (14848 kept out of 15228, 10% discarded) were re-clustered with *dims = 1:20* parameter at resolutions 0.3 and 0.9. Clustering at resolution 0.3 with 14 cell groups was selected as a base for further study. Based on SingleR annotations, cluster 15 from resolution 0.9 were re-introduced in the final dataset as cluster 15, resulting in 15 final cell groups (**Figure S8**). Clusters were finally sorted by size and renamed starting with 1.

### Data analysis

For marker analysis, raw expression data (“RNA” assay) was normalized using Seurat *NormalizeData* function with parameters *normalization.method = “LogNormalize “, scale.factor = 10000.* Afterwards, data was scaled for all genes using Seurat *ScaleData* function with default parameters. Cluster markers of the final dataset were then identified by Seurat *FindAllMarkers* function as described above. GO term enrichment analysis of statistically significant (p-value adjusted < 0.05) cluster markers was performed with the help of clusterProfiler R package [55]. *compareCluster* function was used to compare marker sets per cluster to org.Mm.eg.db database v.3.10 [56] with parameters *fun =“enrichGO”, ont = “BP”, pAdjustMethod = “BH”, pvalueCutoff = 0.05, qvalueCutoff = 0.10, OrgDb = “org.Mm.eg.db”, keyType = “SYMBOL”.*

### Differential abundance testing

In order to find clusters of interest differential abundance testing was performed as described in Chapter 14 of [57] using R edgeR package [58]. Input for the described workflow was produced as follows: cells were assigned a “day” parameter based on the stage of the experiment they were harvested. Abundance matrix of cell counts per sample (columns) per cluster (rows) was produced and used to create a DGEList function output object. Design matrix was produced by model.matrix function from sample IDs by using the “day” parameter as a blocking factor. Clusters to be significantly increased at D10 were cluster 3 (adjusted p=0.005), 5 (adjusted p=0.005) and 15 (adjusted p=0.046) and decreased were cluster 2 (adjusted p=0.025), 6 (adjusted p=0.046) and 10 (adjusted p=0.046), Clusters 3 and 5 were also found to have a significantly lower fraction of control samples compared to the average of others (p-value adjusted 0.018 and 0.018 respectively), as well as to just D10 samples (p-value adjusted 0.002 and 0.002) (**Figure 2B**).

### Inference of pairwise cell interactions

In order to predict pairwise immune cell type interactions, expression matrices of annotated cell types were extracted from each time point of the experiment data. The gene expression data was transformed using the *x*/(*sum*(*x*)\**10000*) formula. Data was then analysed according to the standard CellphoneDB v. 2.1.7 pipeline using *cellphonedb method statistical_analysis* mode with default parameters against v. 2.0.0 database [59]. To match the database, genes were preliminarily renamed to human homologs via R biomaRt v. 2.48.2 package *getLDS* function accessing ENSEMBL *H. sapiens* and *M. musculus* databases [60]. Protein-protein interaction networks of genes forming significant pairs were built and analysed using Search Tool for the Retrieval of Interacting Genes/Proteins (STRING) database v. 11.5 [61] interface (https://string-db.org).

### Data visualization

tSNE scatterplots were made using Seurat *DimPlot* function with parameters *reduction = “tsne”* and *pt.size = 0.2.* Gene expression violin plots were made using Seurat *VlnPlot* function with parameter *pt.size = 0.* Cell type density plots, stackplots and dot plots were made with the help of ggplot2 R package v.3.3.0 [62]. Density plots were built using the tSNE coordinates of each cell per time point with the help of *stat_density_2d* function with parameters *geom = “polygon”, bins = 7, na.rm = T.* Enrichment assay visualization was made by using *dotplot* function from clusterProfiler package with default parameters. Cell composition barplots were created using Microsoft Excel 2019. Gene expression dot plots were created using Seurat R package *DotPlot* function with default parameters. Images were edited using Inkscape v.0.92.3 [63].

## Author contributions

GB, JW designed the study. TY, JW, AK, IB, TB, DST conducted the experimental work and analyzed the data. JW, TB, AK, IB performed parasitic infection in mice, collected samples, isolated and purified hepatic T cells. CAAR isolated and purified the *E. multilocularis* eggs for the infection. GB, DS and JW designed and supervised the experiments in the presented study. TY, JW, GB wrote the original draft. GB, BG, DS, DST contributed to the revised manuscript.

---

**Figure S1.**
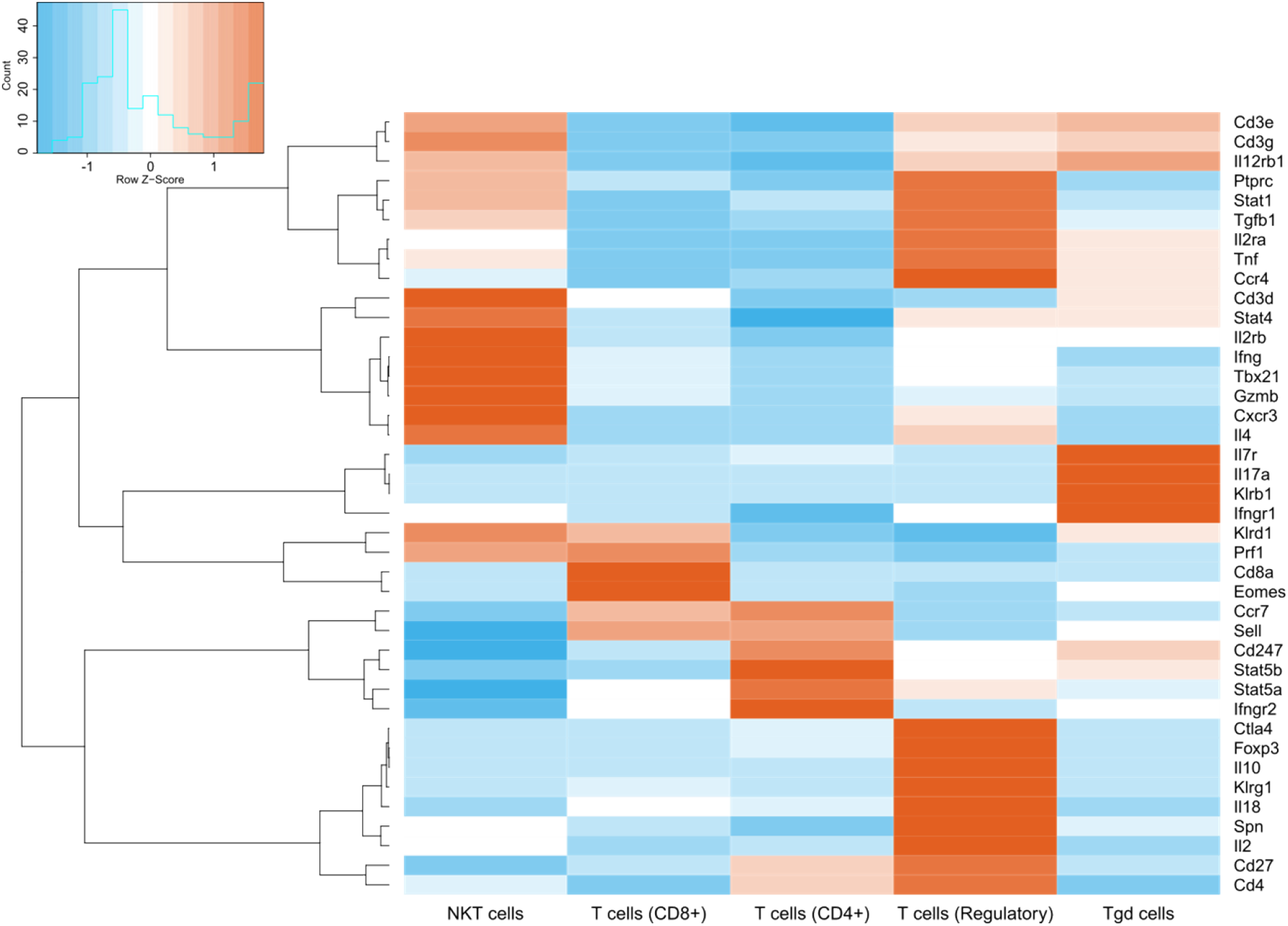
SingleR annotation of the dataset is supported by cell type-defining marker expression. For each SingleR-annotated cell type, the average expression of manually selected cell markers is shown.

**Figure S2.**
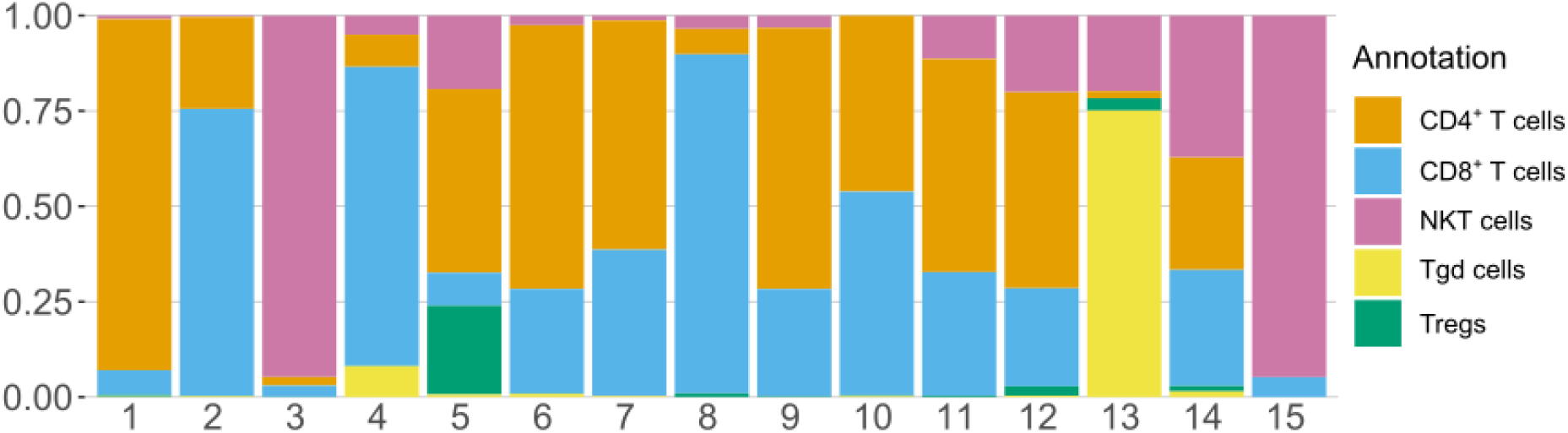
Majority of dataset’s NKT cells is contained within clusters 3 and 15, while most of the Tregs are in cluster 5. For each cluster, the proportion of different cell types is shown.

**Figure S3.**
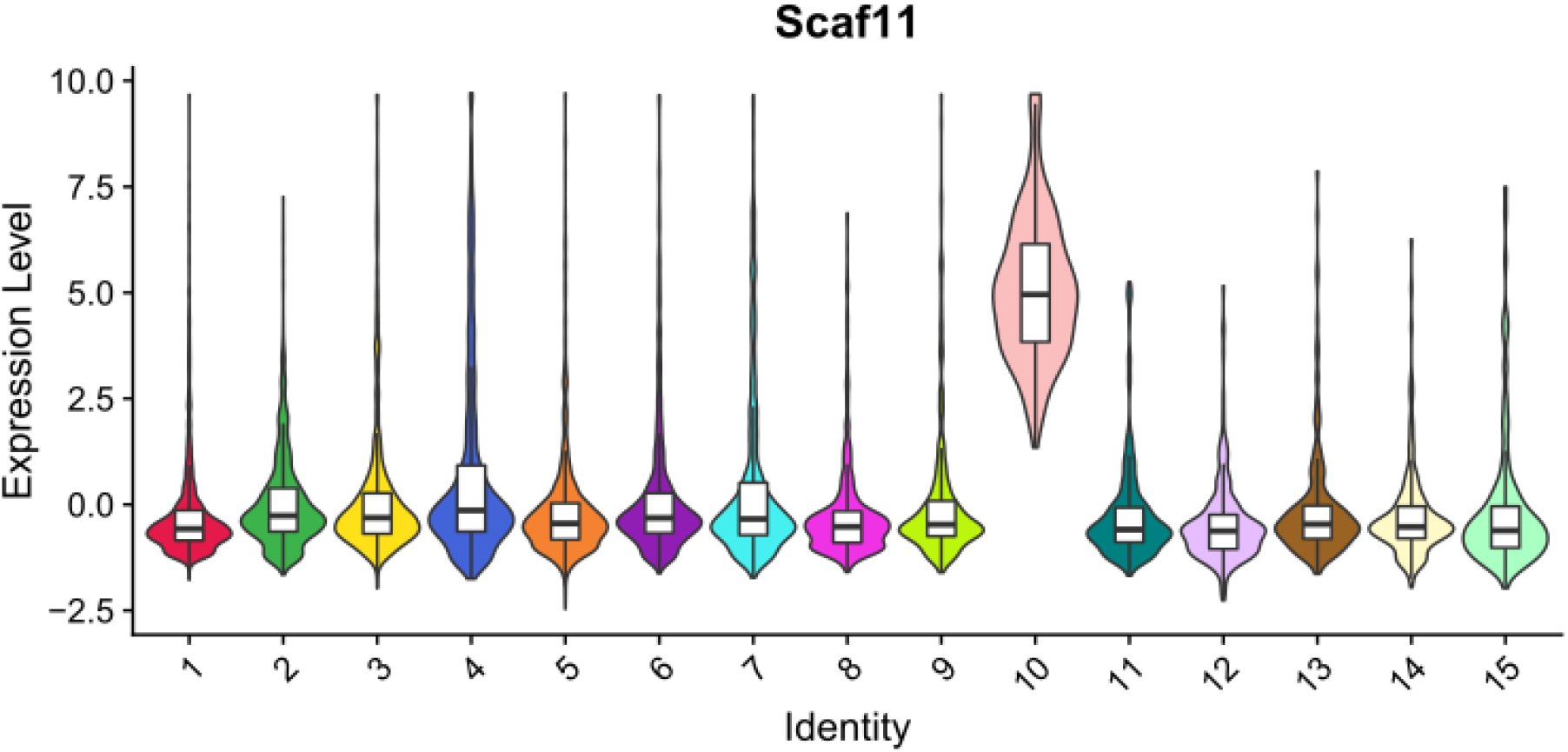
Scaf11 is a strong marker of CD4^+^/CD8^+^ cell cluster 10. Violin plots of expression of Scaf11 in each of the clusters of the datasets are shown, along with a boxplot showing the median and upper/lower quartiles. These results indicate that decrease of a population CD4^+^ and CD8^+^ cells may be the result of pyroptosis.

**Figure S4.**
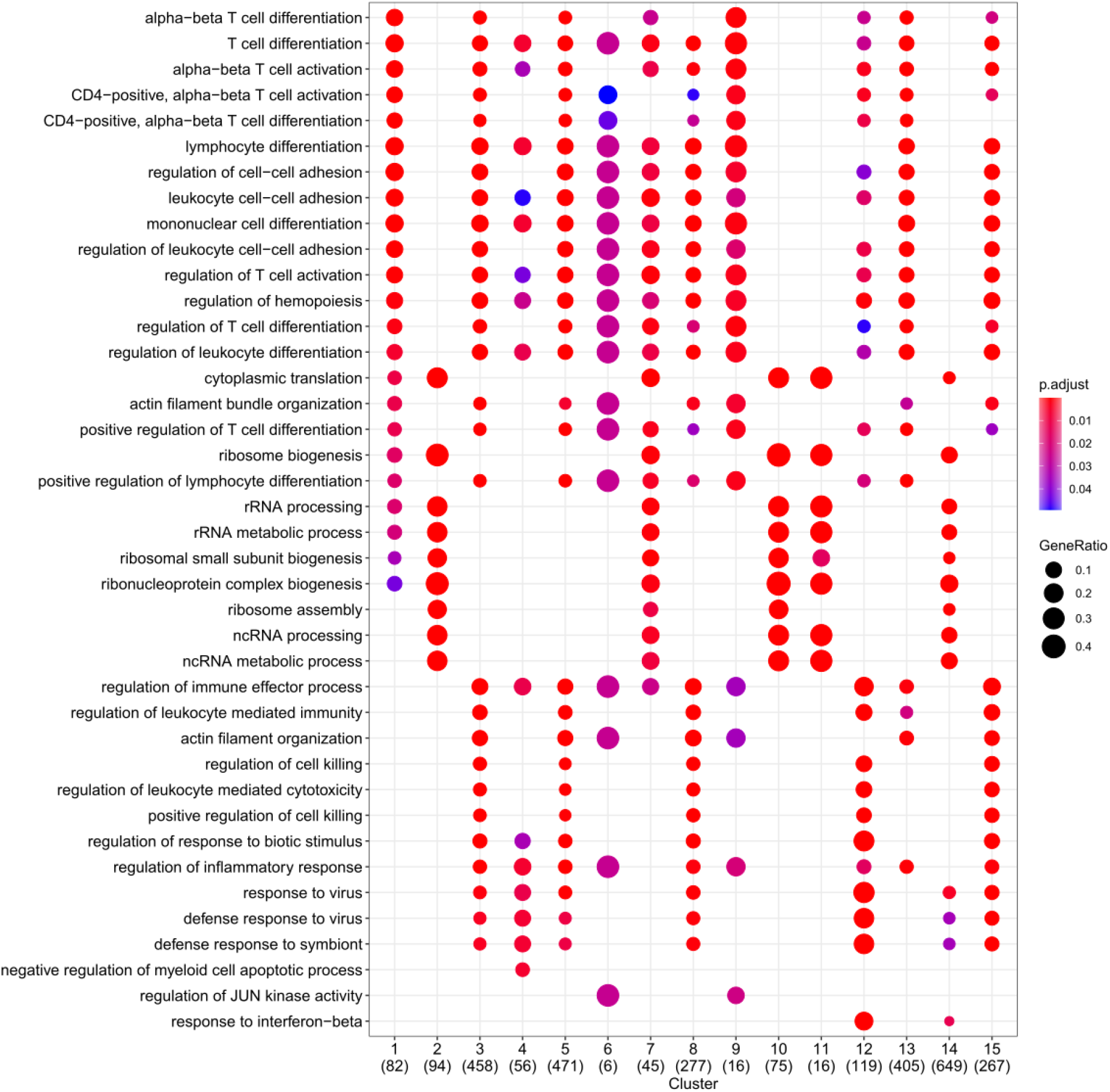
The results of marker set enrichment assay highlight few clusters with ribosomal gene expression on a background of majority immune response-related sets. Enriched pathways are shown on the left, analysed clusters with the number of significant markers (p-value adjusted < 0.05) in brackets on bottom. This indicated that the downregulation of the CD4^+^ and CD8^+^ cell subsets (cluster 2, 7, 10, 11 and 14) may be actively regulated in addition to the induction of pyroptosis in cluster 10

**Figure S5.**
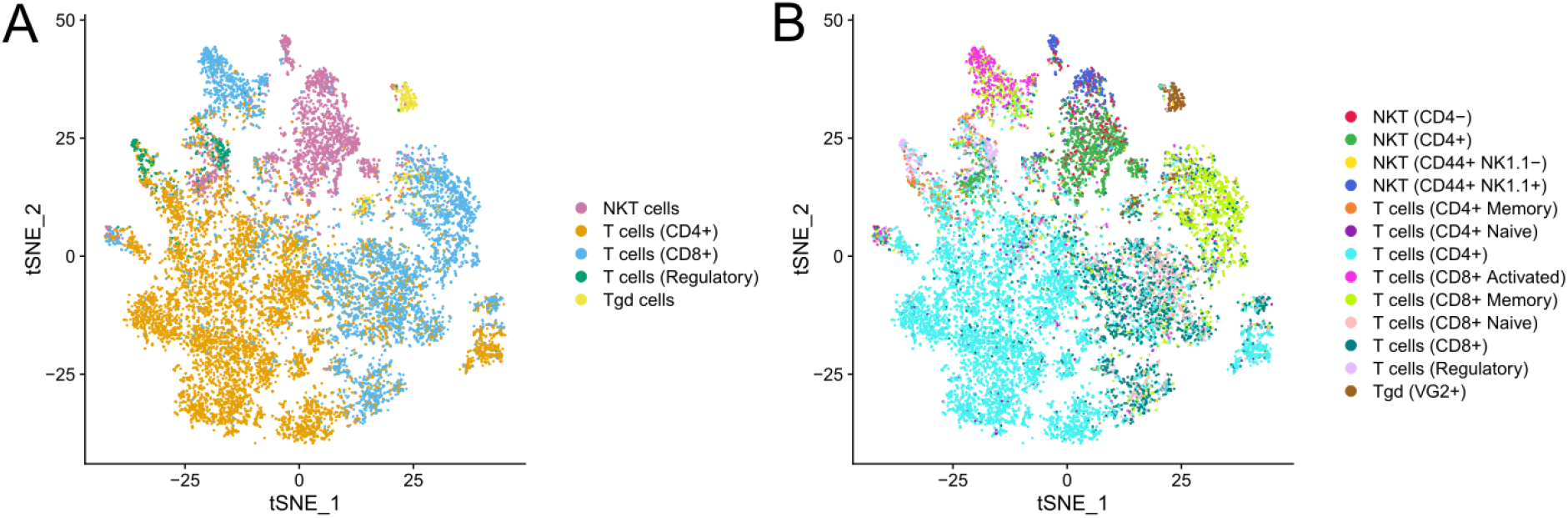
Distribution of T cell subtypes in the experiment. SingleR-based generic cell types (**A**) as well as advanced annotations (**B**) are shown on the tSNE plots representing the data. For exact numbers see **Table S1**.

**Figure S6.**
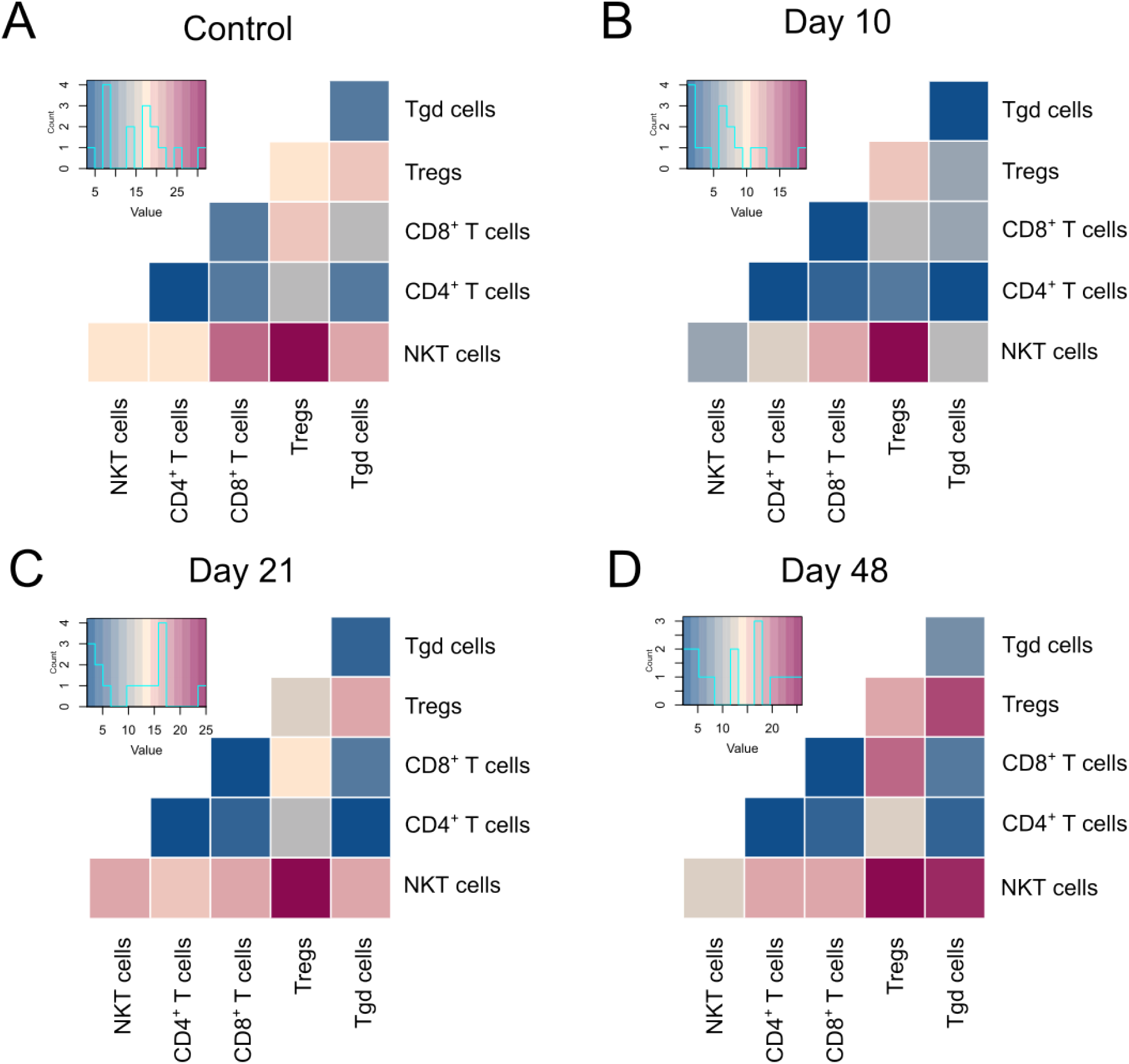
Maps of intercell interactions show a consistently high number of NKT cell and regulatory T cell interactions over the course of the experiment. Heatmaps summarizing statistically significant ligand-receptor interactions per general cell type pair can be seen for control (**A**), D10 (**B**), D21 (**C**), and D48 samples (**D**).

**Figure S7.**
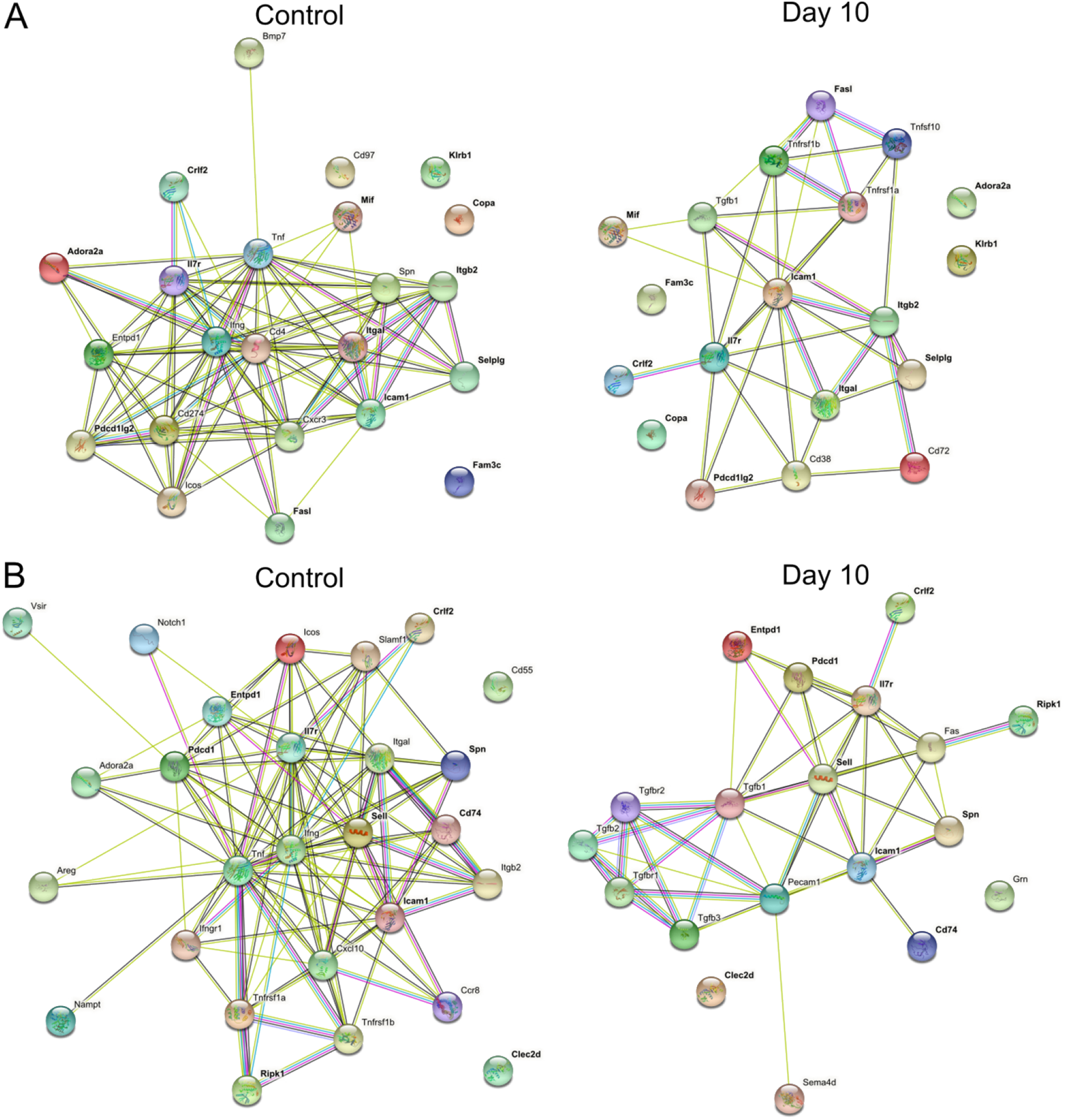
Proteins that form interaction pairs between NKT cells and Tregs display high connectivity within respective cell types. Protein-protein interaction networks are shown for NKT cells (A) and Tregs (B) for genes that form statistically significant predicted NKT-Tregs interaction pairs in control and D10 samples (Figure 4). Proteins that are found both in control and D10 interaction pairs are highlighted in bold.

**Figure S8.**
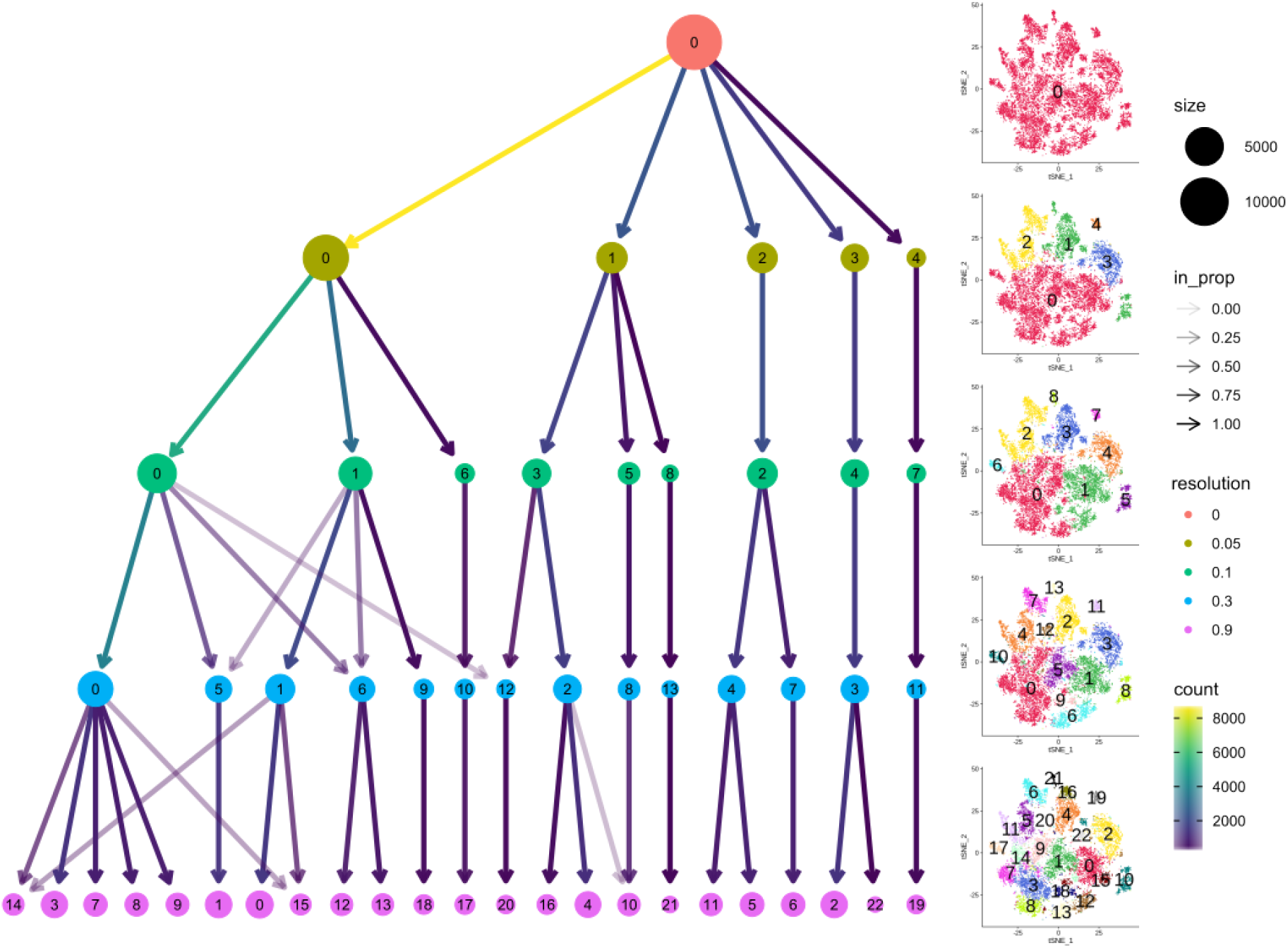
Increasing the clustering resolution allows to highlight a cluster of potential interest. Separation of data on increasing clustering resolution levels is shown schematically (left) as well as on tSNE representations of the dataset (right). Cluster 15 from resolution level 0.9 was reintroduced to the dataset.

